# Intracranial EEG spectral feature analysis and focal brain stimulation reveal affective specialization within dorsal anterior cingulate cortex

**DOI:** 10.1101/2022.03.01.482354

**Authors:** Brian A. Metzger, Prathik Kalva, Madaline M. Mocchi, Brian Cui, Joshua A. Adkinson, Zhengjia Wang, Raissa Mathura, Jay Gavvala, Vaishnav Krishnan, Lu Lin, Atul Maheshwari, Ben Shofty, Sameer A. Sheth, Kelly R. Bijanki

## Abstract

Emotion is represented in several limbic and prefrontal cortical brain areas herein referred to as the Affective Salience Network (ASN). Within this network, less is known about how valence and intensity are processed in the dorsal anterior cingulate (dACC), and how affective processes in dACC compare to activity in other nodes within the ASN. Using a novel spectral feature approach to analyze intracranial electrophysiological data, we discover hemispheric specialization in the dACC such that the right hemisphere is sensitive to intensity while the left hemisphere is sensitive to valence and negative affective bias. We further applied 130 Hz continuous stimulation to the anterior cingulum bundle while patients viewed emotional faces. Faces were rated happier in all patients, an effect modulated by baseline affective bias, suggesting a causal role for the dACC during the processing of external affective stimuli.

## Introduction

The ability to interpret emotional content is an important aspect of social interaction. Failures in this ability are often associated with severe consequences for social communication and are linked to neuropsychiatric disorders including depression. External affective stimuli can be examined in terms of both valence (positive vs. negative) and intensity (subtle vs. overt), where the resulting interpretation of a stimulus and its ecological importance or ‘salience’ are likely influenced by both. Distinct functional networks have been defined for processing affective^1^ and salient^2,3^ stimuli, however, there is overlap in terms of brain anatomy and function in processing of socio-emotional stimuli in these networks.

Brain areas that are known to respond to such stimuli can be broadly categorized as the affective-salience network (ASN). Regions of the ASN include the amygdala^4,5,6^, dorsal anterior and ventral anterior insula (daINS and vaINS^3,6,7,8^), ventral-medial and medial-orbital prefrontal cortices (vmPFC^9^), and dorsal anterior cingulate cortices (dACC^3,10,11,12^). Dysfunctional activity and structural irregularities in regions of the ASN have been linked to social-emotional dysregulation in disorders such as depression^13,14^, and the ASN has been implicated in neural responses modulated by deep brain stimulation into the subcallosal cingulate (SCC) and ventral capsule/ventral striatum (VCVS), white matter fiber pathways innervating the ASN^15,16,17^. The field of affective neuroscience is now critically involved in building our understanding of how affective brain networks dissociably respond to valence and intensity. A better understanding of these functions could help inform the network basis of mood disorders and help produce better-targeted therapies.

Reviewed in Guillory and Bujarski (2014), much is known about affective processing in the ASN, particularly in the amygdala, ventral-medial prefrontal cortex, and the insula. However, much less is known about the role of dorsal anterior cingulate (dACC) for processing emotion. In the current study, we introduce a novel spectral feature detection procedure, and use it to select and analyze intracranial electrophysiological recordings (iEEG) in dACC. Activity in dACC is then compared to multiple other nodes of the ASN (AMY, daINS, vaINS, and vmPFC). We use an Affective Bias emotional evaluation task (ABT) that affords the opportunity to examine the relationship between brain activity and behavior as a function of two constructs that are often conflated in the literature: valence and intensity, plus an additional construct known as negative affective bias, which is the phenomenon whereby external emotional stimuli are interpreted in a manner consistent with one’s own emotional state. Negative affective bias (emotional stimuli interpreted as more negative) has been associated with depression^18–22^, reliably dissociates mood groups, predicts depression treatment responses based on behavioral rating of happy and sad faces^4,5,18,21,23^, and allows us to dissociate emotional valence and intensity.

These advances and improvements in experimental design allow us to investigate the following questions: 1) Is activity in dACC selective for valence (happy vs. sad faces) or intensity (subtle vs. overt emotional expressions)? 2) Can we use neural activity to predict subjective ratings during the Affective Bias task, and hence identify candidate areas for closed-loop brain stimulation to treat affective mood disorders? 3) How does the processing of valence, intensity, and affective bias in dACC relate to the rest of the ASN?

## Results

We collected iEEG data from 16 patients with medication-refractory epilepsy undergoing intracranial EEG monitoring while they participated in the Affective Bias task in which happy and sad faces are rated (Fig. 1A and 1B). Our novel feature detection approach selected prominent features in 228 channels across 5 regions of interest (AMY, dACC, daINS, vaINS, and vmPFC; Fig. 1C).

**Fig. 1.**
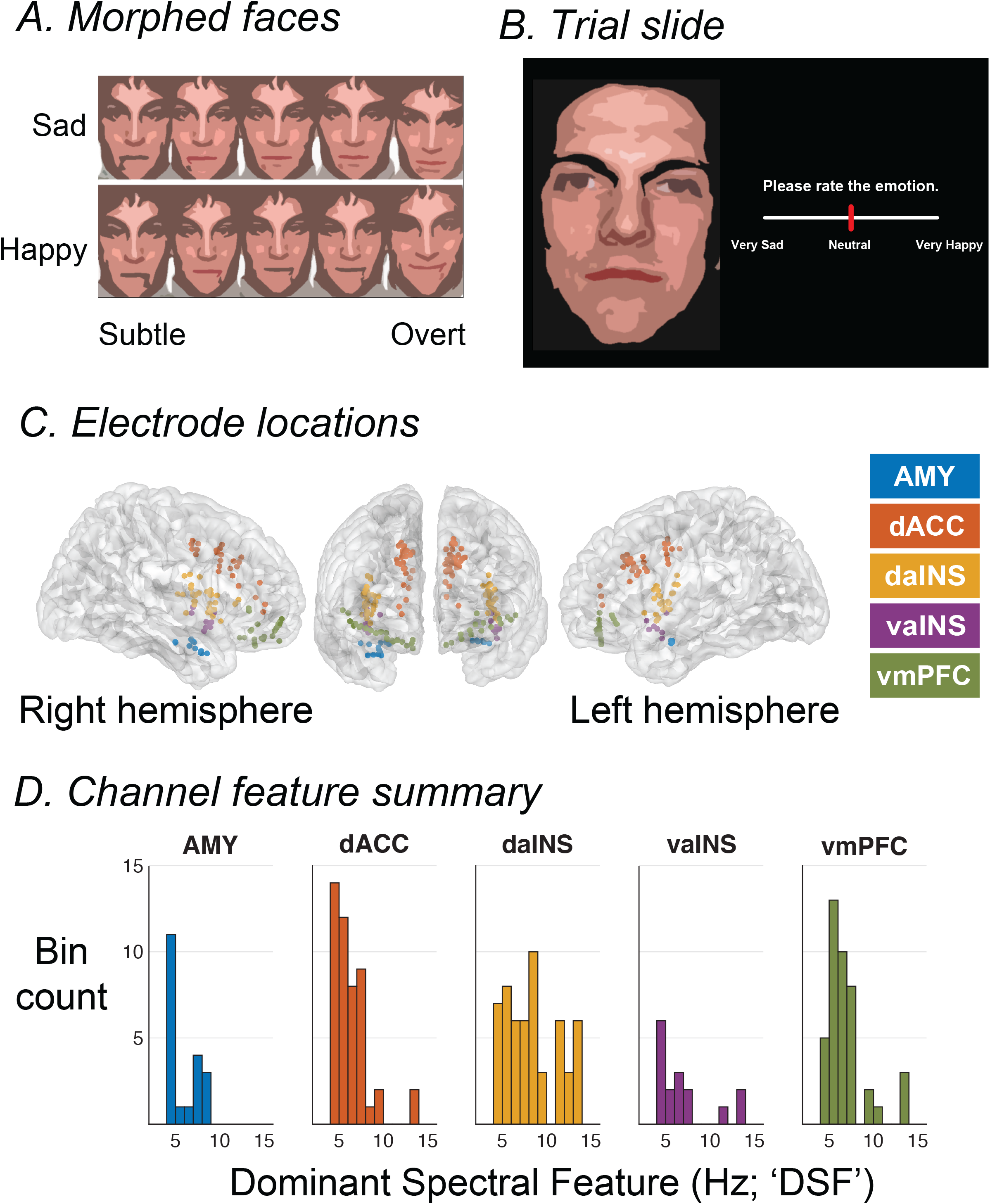
Task and channel recording information. **a** Faces were morphed from neutral to happy, and neutral to sad in steps of 10, 30, 50, and 100%. **b** Faces were presented on a black background. Faces depicted in **a** and **b** are computer generated artistic representations of the actual stimuli, which were presented in their original format (static photographs) during all experiments. The stimuli Participants used a slider bar to rate the emotion of the face. **c** Recording electrodes are colored according to 1 of 5 ROIs. **d** Distribution of dominant detected frequencies (‘Dominant Spectral Feature’) for each ROI.

### Empirical Mode Oscillation Detection (EMOD)

In practice, empirical mode decomposition can be applied to any discretized ordered data, for instance time-series data, or power spectral density (PSD, ordered by frequency). Our approach applies empirical mode decomposition to a vector of spectral power at each wavelet frequency. The output is a series of intrinsic mode functions and the signal residue (or residual). Applied correctly, the residue conforms to a strictly monotonic shape without any local extrema. For any data that conforms to the 1/f power law, the residue is a log-linear 1/f function.

Our approach provides a data-driven method for estimating and removing the background 1/f signal without having to estimate it using linear modeling/regression approaches. Once the 1/f is removed, a dominant spectral feature (DSF) can be detected by simply searching for global and/or local extrema. DSF was identified for each channel separately and power analyses were conducted using a frequency band +/- 2 Hz from the DSF. For example, a DSF at 6 Hz yields a power analysis frequency band of 4-8 Hz. DSF distributions for each ROI are displayed in Fig. 1D. DSFs within the amygdala range from 4-8 Hz and consist of two dominant modes (4 and 7 Hz). DSFs within dACC range from 4-14 Hz and consist of a single peak at 4-7 Hz. Within the anterior insula DSFs ranged from 4-14 Hz with a peak at 4 Hz in ventral channels, while dorsal DSFs conformed to a uniform distribution. DSFs in vmPFC also ranged from 4-14 Hz and consisted of a single dominant mode at 5-7 Hz.

To validate the approach, we compared the goodness-of-fit between EMOD and conventional approaches that estimate 1/f by fitting log-transformed spectral data to a linear function. We measured goodness of fit (r-squared, implemented in MATLAB using ‘robustfit’) for each method and then compared measures using a paired-sample studentized t-test. R-squared values for EMOD ranged from 0.7513 to 0.9998, (mean = 0.9616, std = 0.0306), while r-squared values using a conventional approach ranged from 0.6250 to 0.9943 (mean = 0.8830, std = 0.0672). Across electrodes, our method was associated with stronger fits (mean differences = 0.0787, *t*(227) = 23.47, *p* = 10^-61^). Of all 228 contacts tested, our approach produced a better goodness of fit for 217 out of 228 contacts (95.18%).

### Dorsal anterior cingulate cortex: Spectral power analysis

Our first goal was to determine whether DSF activity varies as a function of either intensity (i.e., subtle vs. overt) or valence (i.e., happy vs. sad). DSF spectral power for each channel was entered into a linear mixed-effects model (LME) that included fixed-effects of intensity, valence, and hemisphere, as well as random effects of participant and channel nested within participant (model notation and summary in Table 1, data visualizations in Fig. 2B and 2C).

**Fig. 2.**
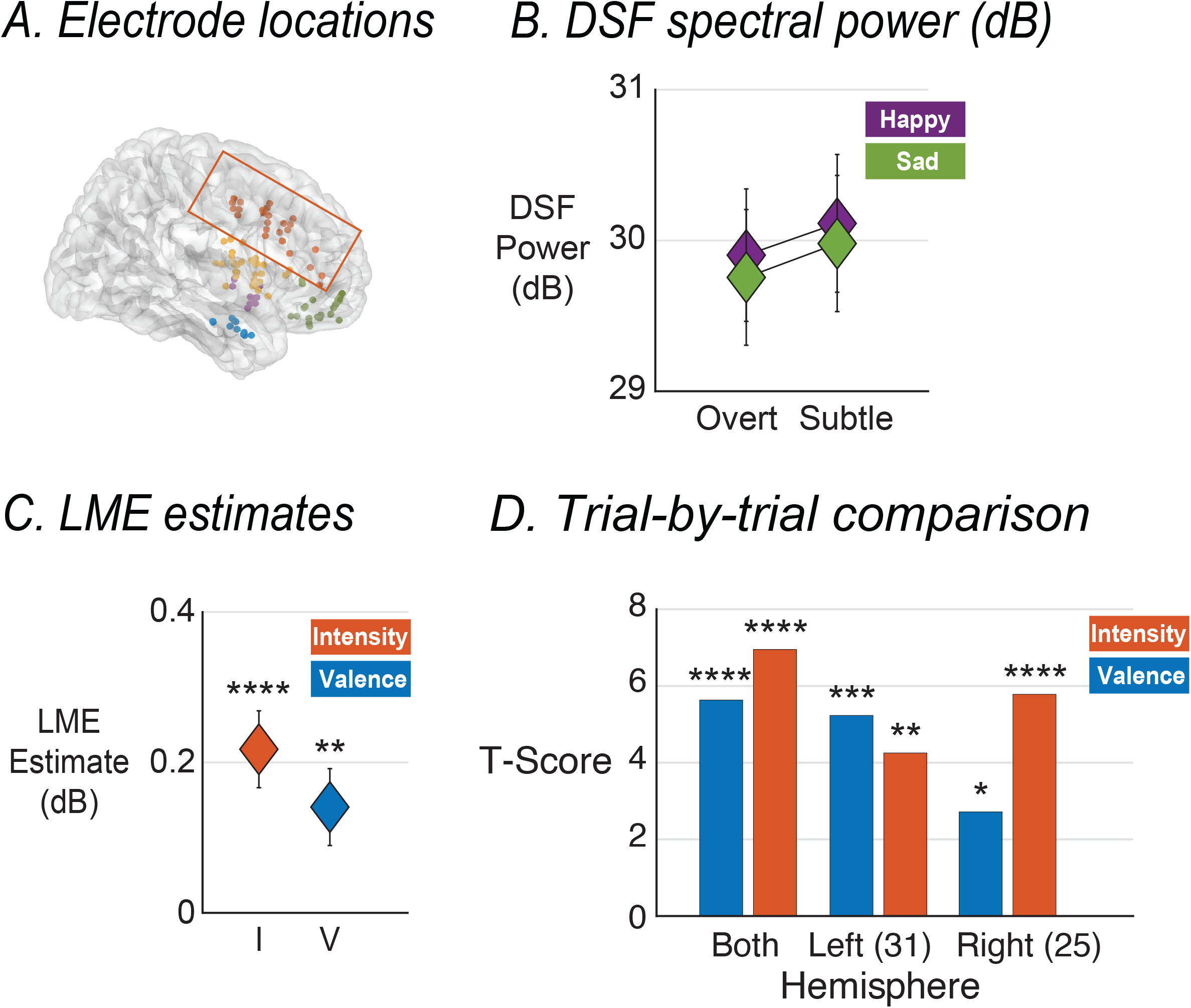
Dorsal ACC: Intensity vs. Valence. **a** Electrode locations colored according to ROI. dACC contacts are indicated by the orange box. **b** Mean spectral power (dB) shown as a function of intensity (‘O’ = Overt, ‘S’ = Subtle) and valence (Happy vs. Sad) averaged across dACC channels. **c** Fixed-effect estimates (dB) for Intensity (I) and Valence (V) from a linear mixed-effects model. **d** Comparisons of trial-by-trial correlations between spectral power and either intensity-coded ratings (orange bars) or valence-coded ratings (blue bars) displayed as a t-score (All error bars +/− 1SEM; * = < 0.05, ** < 0.01, *** < 0.001, **** < 0.0001).

**Table 1.** Linear mixed-effects summary.

| ROI | Model Coefficients | Fixed Effect | Estimate | P-Value |
| --- | --- | --- | --- | --- |
| Amygdala | I + V + H + (1\|SubID/ChanNum) | Intensity | 0.1011 | 0.0362 * |
|  |  | Valence | -0.451 | 0.0003 *** |
|  |  | Hemisphere | -0.188 | 0.6099 |
| Dorsal ant. cingulate | I + V + H + (1\|SubID/ChanNum) | Intensity | 0.2176 | <0.0001 **** |
|  |  | Valence | 0.1407 | 0.0066 ** |
|  |  | Hemisphere | -0.6470 | 0.4691 |
| Dorsal ant. insula | I + V + H + (1\|SubID/ChanNum) | Intensity | 0.1436 | <0.0001 **** |
|  |  | Valence | 0.1115 | 0.0022 ** |
|  |  | Hemisphere | 0.5804 | 0.4768 |
| Ventral ant. insula | I + V + H + (1\|SubID/ChanNum) | Intensity | 0.0338 | 0.7100 |
|  |  | Valence | 0.1220 | 0.1820 |
|  |  | Hemisphere | -0.4997 | 0.7270 |
| Ventral med. PFC | I + V + H + (1\|SubID/ChanNum) | Intensity | 0.3134 | <0.0001 **** |
|  |  | Valence | 0.0951 | 0.0194 * |
|  |  | Hemisphere | -3.2598 | 0.0191 * |

Statistical significance: \* < 0.05, \*\* < 0.01, \*\*\* < 0.001, \*\*\*\* < 0.0001
Estimates reflect the modeled difference between factors within each fixed effect. Intensity estimates reflect Subtle>Overt; positive values reflect greater DSF power for Subtle faces. Valence estimates reflect Sad>Happy; positive values reflect greater DSF power for Sad faces. Hemisphere estimates reflect Right>Left; positive values indicate greater DSF power for Right hemisphere contacts.

The LME resulted in significant fixed-effects for valence such that sad faces evoked reduced DSF activity relative to happy faces in dACC (fixed-effect estimate = 0.1407, *p* < 0.0001), and for intensity such that overt faces evoked reduced DSF activity relative to subtle faces (fixed-effect estimate = 0.2176, *p* < 0.0001). No statistically significant differences were observed between hemispheres (*p* = 0.4691).

### Trial-by-trial correlations: Predicting valence vs. intensity

LME results suggest that dACC is sensitive to emotional valence and intensity, albeit with larger effects for intensity. We next sought to determine whether DSF activity was predictive of task behavior. Emotional ratings provided by patients were correlated with DSF activity at the trial level. Correlations were run using two methods of calculating the emotional rating. Ratings were initially recorded along a valence scale ranging from 0 (‘Very Sad’) to 1 (‘Very Happy’) with ‘Neutral’ falling at the midpoint (0.5). These ratings were also transformed to an intensity scale by subtracting sad face ratings from 1. The consequence of this transformation is that maximally happy face ratings and maximally sad face ratings are equal in magnitude. We calculated correlation coefficients separately for each scale. Significance at the level of the ROI was determined by conducting a t-test of the Fisher’s Z-transformed correlation coefficients against 0. ROI t-tests were conducted by first including all channels in the analysis regardless of hemisphere, and then repeated for contacts in each hemisphere separately. The resulting *p*-values were corrected using a false discovery rate of 0.05 (Supplementary Table 1).

DSF spectral power was predictive of valence-coded emotional ratings in the dorsal ACC (Fig. 2D). Dorsal ACC contacts showed a positive correlation between spectral power and valence-coded ratings such that an increase in DSF spectral activity was associated with happier ratings. Within dACC, valence-coded correlation coefficients in the left hemisphere were larger and more consistent (*rho*_mean_ = 0.09, *p* <0.001) compared to the right hemisphere (*rho*_mean_ = 0.05, *p* <0.012).

DSF spectral power was also predictive of intensity-coded emotional ratings in dACC (Fig. 2D). Correlation coefficients were negative such that an increase in DSF spectral power was associated with a decrease in intensity ratings. Separating dACC by hemisphere revealed stronger intensity-coded predictive associations in the right hemisphere (*rho*_mean_ = −0.10, *p* <0.001) compared to the left hemisphere (*rho*_mean_ = −0.07, *p* = 0.001).

Taken together, the LME and trial-by-trial correlation analyses suggest an interesting double dissociation in dACC. DSF spectral activity was more predictive of valence in the left hemisphere, while more predictive of intensity in the right hemisphere. Furthermore, the direction of the predictive associations went in opposing directions. Valence was positively correlated such that happier ratings were associated with greater DSF activity, while intensity was negatively correlated such that more intense ratings were associated with reduced DSF activity. Areas within the affective salience network are not only more sensitive to the intensity of the emotion (i.e., subtle vs. overt expressions) compared to valence (i.e., happy vs. sad, positive vs. negative), but are also uniquely predictive for intensity.

### Predicting Affective Bias

To better understand the relationship between affective bias and Affective Salience Network iEEG activity, we transformed the emotional ratings from the behavioral task into affective bias scores by subtracting the observed rating from an expected rating, and then correlated these scores with DSF spectral power on a trial-by-trial basis.

Expected ratings came from a separate study consisting of participants recruited via Amazon’s Mechanical Turk (MTurk). MTurk participants were initially screened for depression symptoms (n = 400). The face rating task was only administered to participants exhibiting no depression symptoms (n= 86). Averaged “expected” values were calculated for each stimulus (Extended Data Fig. 1a).

At the ROI level (Fig. 3B), affective bias scores were correlated with DSF spectral power only in dACC, and within dACC, correlation coefficients were larger and more significant in left hemisphere contacts (*rho*_mean_ = 0.06, *p* = 0.002) relative to right hemisphere contacts (*rho*_mean_ = 0.04, *p* = 0.081). Correlations between affective bias scores and DSF spectral power were positive, suggesting that lower spectral power in dACC is associated with increased negative bias (interpreting faces as sadder than ratings from healthy controls; such interpretation is an indicator of negative mood state).

**Fig. 3.**
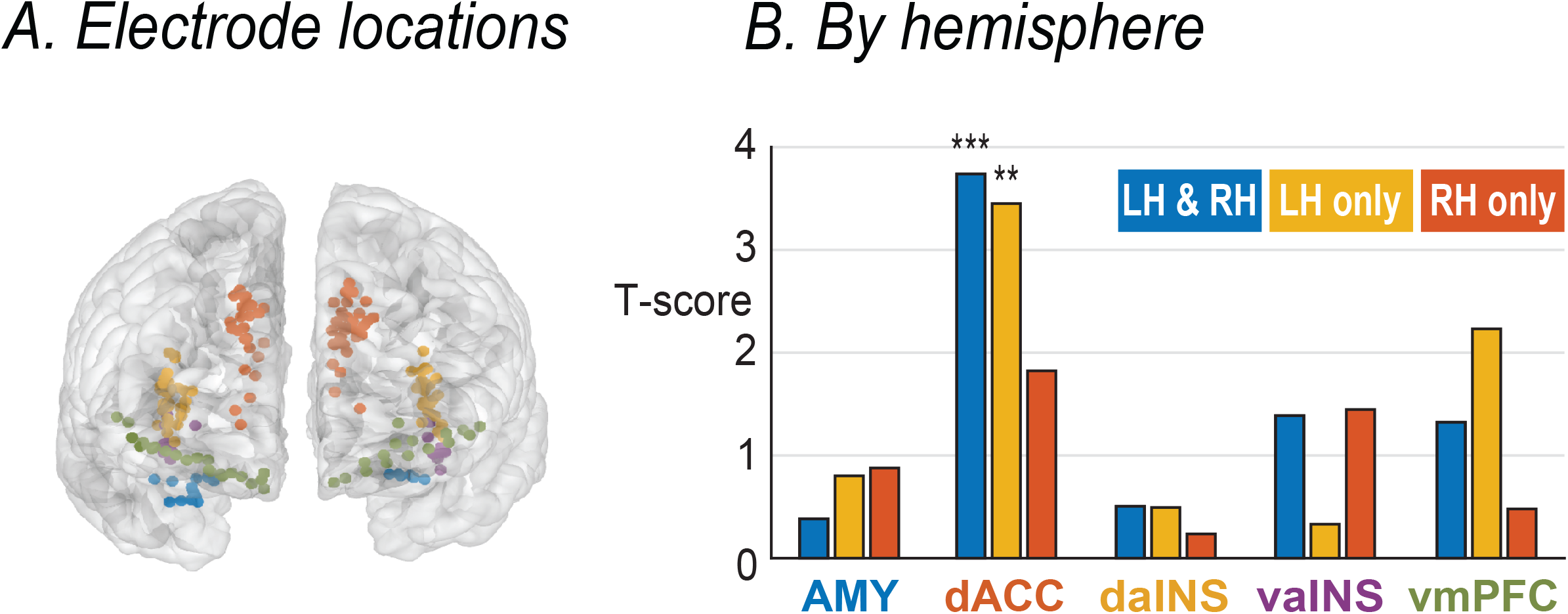
Predicting DSF spectral power from Affective Bias (expected-observed emotional ratings). **a** Electrode locations colored according to ROI. **b** ROI-level comparisons of trial-by-trial correlations between Bias score (expected-observed emotional ratings) displayed as t-scores for each ROI (blue bars = contacts from both hemispheres included, orange bars = only right hemisphere contacts, gold bars = only left hemisphere contacts; ** < 0.01, *** < 0.001).

### Stimulation of ACC during Affective Bias

The association between activity in dACC and negative affective bias demonstrates an important role for the dACC in the evaluation of external emotional stimuli. To test the causality of this role, we conducted an additional experiment in which 4 patients (3 of whom were included in the intracranial EEG analyses described above) received 130 Hz continuous stimulation to the anterior cingulum bundle during the Affective Bias task. Stimulation runs were blinded from patients and sham controlled. Stimulation locations are displayed in Fig. 4A. One patient (‘YDX’) received bipolar stimulation, while the other 3 received monopolar stimulation.

**Fig. 4.**
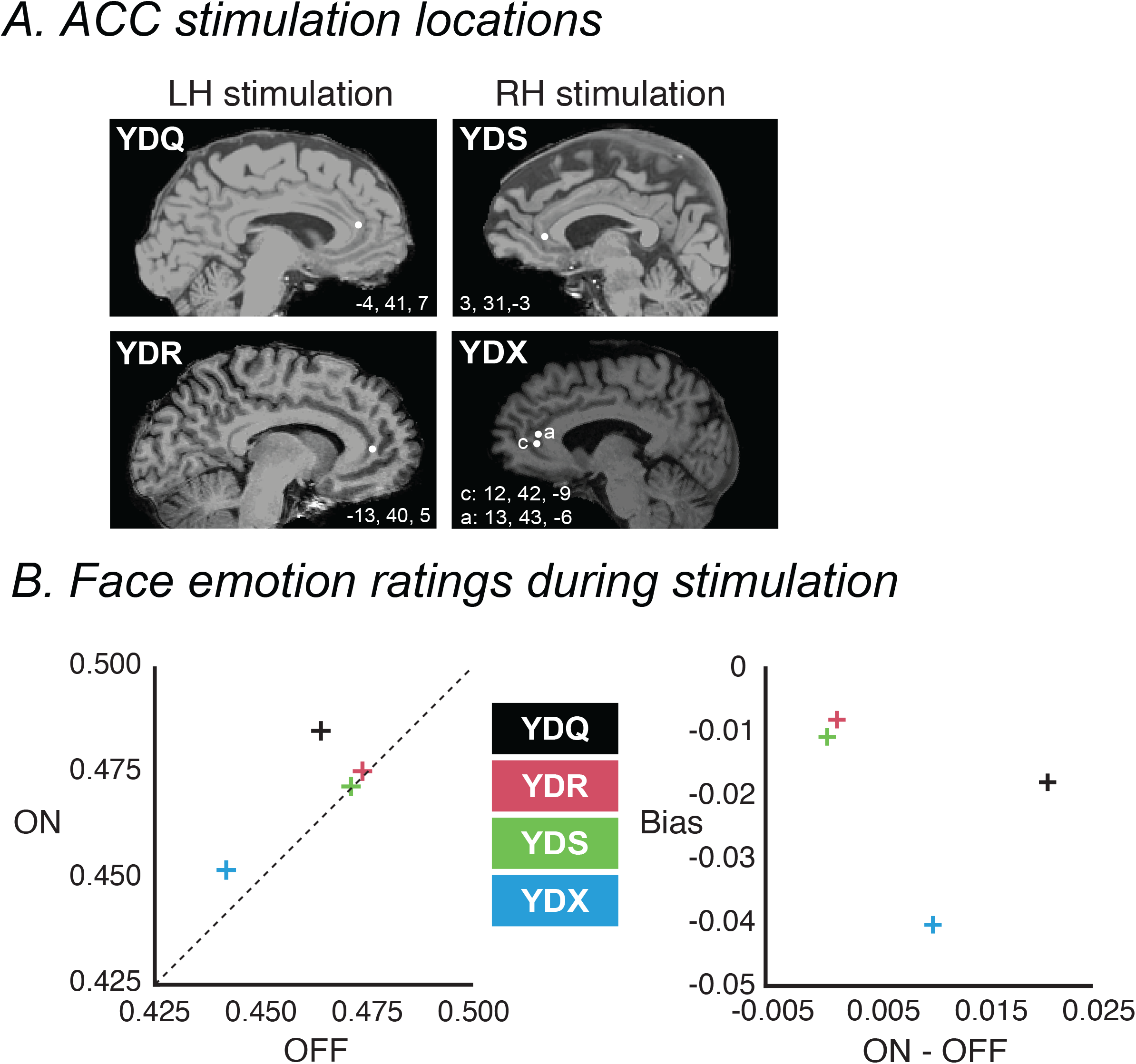
Task performance during stimulation to ACC. **a** ACC stimulation locations displayed for each subject. YDQ and YDR received left hemisphere stimulation, YDS and YDX received right hemisphere sitmulation. **b** Left panel: emotional ratings for each patient as a function of whether stimulation was off (x-axis) or on (y-axis). Dashed line represents line of equivalence. Ratings above the dashed line indicate ratings were higher while sitmulation was on. **b** Right panel: difference between stimulation on and stimulation off (x-axis) compared against average Affective Bias score (expected-observed). Subjects with low AB show little to no change in task ratings as a function of stimulation.

Average ratings were higher during stimulation compared to no stimulation in all four patients. As shown in Fig. 4B (left), two patients showed large differences between stimulation on and off, while the other two showed relatively small effects. To try to explain the variation in stimulation effect size, we tested for a relationship between negative affective bias and stimulation-induced changes in emotional ratings. For instance, if stimulation restores healthy function in dACC, stimulating dACC would only be expected to normalize emotional processes in the unhealthy brain. Stimulating ACC in someone with a high degree of negative bias could be associated with a greater effect of stimulation on task performance compared to someone with a low degree of negative bias. To explore this hypothesis, we compared affective bias scores from the baseline session of the task and the stimulation effect (same-day ratings with stimulation on vs. off). Data were entered into a linear mixed-effect model that included fixed effects of valence, pre-stimulation Bias scores and a random effect of patient. Pre-stimulation baseline bias scores were highly predictive of the stimulation effect (fixed-effect estimate = −0.6154, *p* = 10^-18^; Fig. 4B, right). In other words, patients with a higher degree of negative bias at baseline also have greater stimulation effects. Together, these data suggest a causal role for dACC during the evaluation of external emotional stimuli.

### Behavior

To determine whether ratings of faces with subtle emotional expressions (i.e., neutral and 10% morphed) differ as a function of valence (i.e., happy vs. sad), subtle face data were entered into a mixed-effects model that included a fixed effect of valence (happy vs. sad), and random effects of participant and channel nested within participant. We observed no statistically significant differences between subtle happy and subtle sad faces (*p* = 0.2865). To determine whether the difference between overt faces and subtle faces was stronger for happy relative to sad faces, we calculated the absolute difference between overt and subtle for each patient separately for happy and sad faces and entered these data into the same mixed effects model. We saw no differences as a function of valence (*p* = 0.8787). Extended Data Fig. 1 shows a behavioral summary as a function of valence and intensity for the epilepsy and MTurk cohorts.

### Affective Salience Network: ROI comparisons

We extended the LME and trial-by-trial correlation analyses to four other ROIs of the ASN (AMY, daINS, vaINS, and vmPFC). Comparisons of results across ROIs are displayed in Extended Data Fig. 2 (DSF spectral power LME results) and Extended Data Fig. 3 (trial-by-trial correlation comparisons).

LME modelling revealed significant differences as a function of intensity in all ROIs except for vaINS (fixed-effect estimate = 0.0338, *p* = 0.0338), such that overt faces evoked reduced DSF activity relative to subtle faces. The largest effect was observed in vmPFC (fixed-effect estimate = 0.3134, *p* < 0.0001), followed by dACC (fixed-effect estimate = 0.2176, *p* < 0.0001), then daINS (fixed-effect estimate = 0.1436, *p* < 0.0001), and finally the amygdala (fixed-effect estimate = 0.1011, *p* < 0.0362). Likewise, significant differences as a function of valence were observed in all ROIs except for vaINS (fixed-effect estimate = 0.1220, *p* = 0.1820). Sad faces evoked increased DSF activity relative to Happy faces in the amygdala (fixed-effect estimate = −0.451, *p* = 0.0003), while sad faces evoked reduced activity relative to happy faces in dACC (fixed-effect estimate = 0.1407, *p* < 0.0001), daINS (fixed-effect estimate = 0.1115, *p* = 0.0022), and vmPFC (fixed-effect estimate = 0.0951, *p* = 0.0194). Differences in DSF spectral power as a function of hemisphere were only observed in vmPFC such that DSF spectral power was greater in the left hemisphere (fixed-effect estimate = −3.2598, *p* = 0.0191, all other p-values > 0.05).

Together, the LME show that most ROIs (apart from vaINS) are sensitive to emotional valence and intensity. The amygdala appears to be more sensitive to valence, while dACC, daINS, and vmPFC are more sensitive to intensity.

Trial-by-trial correlation analyses showed that DSF spectral power was predictive of valence-coded emotional ratings in dACC and amygdala (Extended Data Fig. 3). Amygdala contacts showed a negative correlation between spectral power and valence-coded ratings such that an increase in DSF spectral activity was associated with sadder ratings. Within the amygdala, correlation coefficients were larger and more consistent in the right hemisphere (*rho*_mean_ = 0.07, *p* = 0.017) compared to the left hemisphere (*rho*_mean_ = 0.06, *p* = 0.343), however this could be due to low number of left-hemisphere contacts (n = 6, compared to n = 13 in the right hemisphere). DSF spectral power was consistently predictive of intensity-coded emotional ratings of all ROIs except vaINS. ROI-averaged correlation coefficients were largest and most consistent in dACC (*rho*_mean_ = −0.08, *p* <0.001) and vmPFC (*rho*_mean_ = −0.10, *p* <0.001), followed by daINS (*rho*_mean_ = −0.06, *p* <0.001) and finally in the amygdala (*rho*_mean_ = −0.05, *p* = 0.004). For all ROIs, the correlations were negative such that an increase in DSF spectral power was associated with a decrease in more intense ratings, which is consistent with the results from the LME reported in the prior section. Separating ROI contacts by hemisphere revealed stronger predictive associations in the right hemisphere for the amygdala, dorsal ACC, vaINS and vmPFC. Only daINS showed stronger predictive relationships in the left hemisphere.

Taken together, the LME and trial-by-trial correlation analyses suggest that areas within the affective salience network are not only more sensitive to the intensity of the emotion (i.e., subtle vs. overt expressions) compared to valence (i.e., happy vs. sad, positive vs. negative), but are also uniquely predictive for intensity.

## Discussion

Using a novel spectral feature approach to analyze iEEG data, we show hemispheric specialization in dACC. The right hemisphere was sensitive to intensity while the left hemisphere was sensitive to valence and negative affective bias. We further applied 130 Hz continuous stimulation to the anterior cingulum bundle and found the changes in emotional ratings were modulated by baseline affective bias. These data strongly suggest a causal role for dACC during affective processing.

### Targeting positive vs. negative affect

Depression is a highly variable disorder characterized by several potential biotypes, each of which is associated with dysfunction in distinct brain networks^14^. For instance, anhedonia is associated with hyperactivation of vmPFC^24,25^, while rumination is associated with hyperactivation of the default mode network, which includes nodes within dACC. Our LME analysis of activity in dACC revealed strong effects for intensity and valence. We were also reliably able to predict emotional ratings and Bias scores from DSF trial-level activity in dACC, the latter of which was only achieved in dACC. Together, these data suggested that dACC is critical for processing the emotional content of faces. Previous research in the dACC suggests that this area is important for integrating different dimensions of motivationally-significant stimuli to encode value or emotion^26,27^. Our findings further support this view, suggesting dACC may play a role in integrating valence and intensity components of emotion during affective and salience processing. Deficits in affective processing seen in mood disorders such as depression may arise, then, from the inability to integrate multiple components of affective stimuli or update affective perception effectively due to hypo-functional connectivity or hypo-activation^14,28^ of the dACC. Direct electrical stimulation to this region could potentially restore this integrative and adaptive function of the dACC. In fact, our data showed that stimulation to ACC during the Affective Bias Task was able to increase emotional rating of faces in each of 4 patients who received stimulation. Importantly, this effect was specific to those exhibiting negative affective biases, and showed an increasing effect with increasing baseline negative affective bias, which suggests that dACC stimulation is not simply altering affective interpretation of valence (i.e. all patients would exhibit an increase in positive ratings), or intensity (i.e. all patients would exhibit an increase positive and negative ratings) alone, but rather is adjusting integration of the two for those exhibiting deficits, and not surpassing ‘normal’ function. Thus, dACC may serve as an ideal location from which to monitor brain activity to identify states (i.e., moments) of dysfunction, and then stimulate in closed-loop fashion.

### Affective valence vs. intensity

Valence refers to positive vs. negative, while intensity refers to some measurable, graded distance from neutral. Researchers must be careful not to confound them. Contrasting positive vs. neutral, for instance, is a comparison of both valence and intensity. Contrasting positive vs. negative is only a comparison of valence so long as the positive and negative stimuli have been controlled for intensity, which can be difficult. One approach is to compare arbitrarily defined intensities^30,35,36^, or to use an ordinal scale ranging from neutral (least intense) and a given emotion (most intense), however both approaches confound valence and intensity. Another approach is to use an ordinal scale ranging from negative (lowest ranking) to neutral (middle ranking) to positive (high ranking), however this approach, without a transformation of the ratings, also confounds valence with intensity. Our study uses the Affective Bias task, which can assess both valence and intensity using a continuous scale (presented as extremely sad through extremely happy, with neutral exactly between the two). We apply a transformation to the valence-coded ratings that allows us to independently assess valence and intensity. The trial-by-trial correlation analysis strongly favored the intensity-coded ratings in every area, suggesting that the predominant role of the ASN is to detect emotional salience (hence our motivation for coining it the Affective Salience Network).

### Affective subnetworks

Disentangling the region-specific functions of the Affective Salience Network is an essential step towards both characterizing emotional processing in the brain, and effectively treating emotion-related pathologies. Data from scalp EEG^29,30^ and fMRI^31,32^ tend to support the Valence Hypothesis, which proposes that emotion is processed in a valence-specific manner such that the left hemisphere processes positive valence while the right hemisphere processes negative valence. The current data suggest that at the network level, activity within the ASN is more sensitive to intensity compared to valence. For instance, the LME and the trial-by-trial correlation analyses provide strong evidence that several areas within ASN are specifically sensitive to intensity (subtle vs. overt; daINS, vmPFC, left dACC), while some are specifically sensitive to valence (happy vs. sad; amygdala, right dACC), and one region showing mixed responsiveness to both (vaINS). Further, within the ASN, dACC and possibly vaINS show hemispheric specialization such that the left hemisphere contacts are more sensitive to valence while right hemisphere contacts are more sensitive to intensity.

An interesting point of mechanistic integration of the current findings is with the anatomical distribution of Von Economo neurons (VENs)–a population of neurons with a unique bipolar morphology and a spindle-shaped soma, expressed almost exclusively in the human anterior cingulate cortex and anterior insula^33^. These cells are relatively large and are thought to facilitate rapid communication across distant brain regions^33^. in addition to their putative role in integrating disparate information streams to mediate socio-affective functions. Abnormalities in Von Economo neurons tend to translate to regional dysfunction in the aINS and ACC and are related to deficits in social communication such as in autism and frontotemporal dementia^34^. These findings may with further research help characterize a unique means of socio-affective processing in the human brain, reliant on the ASN.

### Comparison of feature detection approaches: EMOD vs. conventional approaches

Conventional approaches of analyzing EEG neural data have relied on restricting analysis to a predefined canonical frequency band (i.e., theta, alpha, etc.); neural activity gets defined as power in an arbitrarily defined oscillatory bands, which are kept the same for analyses over all brain regions for scalp EEG^29,30^ and intracranial EEG^37,38^. However, recent findings cast doubt on this approach showing that the dominant frequency of an oscillation, or neural feature, can vary along an anatomical dimension^39^. Recent approaches have shifted away from frequency-band analyses based in favor of data-driven selection approaches that search for dominant features in power spectral density^39–42^.

Our study introduces a data-driven approach that empirically derives dominant electrophysiological features of interest from spectral power density independently for each analyzed data channel and then uses spectral power in the detected feature. In this way, we also begin to specify oscillatory frequencies of interest more efficiently for future stimulation studies, rather than examining arbitrarily defined broad frequency bands which may or may not be relevant to the region-specific responses of interest. Alternative approaches rely on iterative rounds of linear fitting and signal regression. Fitting is not without error, and as such can distort the central frequency or feature magnitude when estimating feature characteristics from spectral data. Our novel feature detection approach identifies spectral features without the need to first identify and remove the 1/f power law function. In the present data, EMOD reliably detected spectral features and feature magnitudes more accurately compared to conventional approaches. Of the identified 228 channels in 5 ROIs (AMY, dACC, daINS, vaINS, and vmPFC) with significant neural features, 95% were associated with a better goodness-of-fit for EMOD compared to conventional approaches.

Using our method, we found that the dominant feature central frequency varied considerably both within and across ROIs, spanning the entire range of canonical theta-band and alpha-bands (i.e., 4-8 Hz, and 8-14 Hz). Close inspection of Fig. 1D shows little consistency in terms of the central frequency of the detected feature. The distribution of dominant features varies greatly across ROIs in multiple ways. There are differences in terms of range (narrow: AMY; wide: all others), and shape (unimodal: dACC, vaINS, vmPFC; bimodal: AMY; uniform: daINS), suggesting that canonical approaches focusing on pre-defined ranges (i.e., theta or alpha) fail to capture significant and meaningful differences in brain activity for any contrast of interest, particularly if electrodes sampled from a large ROI or across patients are included.

### Future directions

Several open questions remain regarding how to assess emotion precisely but broadly from a neuroscientific perspective. One limitation of the current study concerns the use of emotional faces. The extent to which our findings generalize remains an open question. For instance, it is unknown whether our findings generalize to other domains of emotion both in terms of category (fear, anger, disgust, etc.), expression (facial, vocal), or perception (vocal pitch, image brightness, etc.). Future research should also seek to determine whether the Affective Bias Task can be used to classify or index mood, and whether it can predict treatment outcome and track changes in mood longitudinally. Clinical questionnaires including the Hamilton Depression Scale, Montgomery-Asberg Depression Rating Scale, and the Beck Depression Inventory, are only sensitive to changes in depression over relatively long periods of time. Hence, Affective Bias behavioral tasks can serve as a useful adjunct to these scales especially when the need arises to test the efficacy of brain stimulation on a short time scale. Furthermore, some individuals are incapable of completing a depression inventory, for instance locked-in patients who are incapable of providing a vocal or written response, or for patients who suffer from alexithymia and are incapable of describing their own emotional state.

Several open questions also exist regarding the Affective Salience Network. It remains to be determined whether the ASN would show the same pattern of response for happiness/sadness in terms of valence and intensity for other emotions such as fear, or disgust, or anger. Future research should seek to understand the long-term functional changes within the Affective Salience Network both as a function of changes in mood, as well as a function of brain stimulation.

In summary, the findings from linear mixed-effects modeling and direct intracranial electrical stimulation in this study suggest a causal role for dACC during the processing of emotional content. We have demonstrated the capacity for neural activity in the dACC to predict emotional behavior at the level of individual trials (negative bias in the rating of emotional stimuli). Furthermore, this study demonstrates trial-level predictive associations between baseline affective bias and stimulation effects on perception, suggesting that individuals with greater baseline negative bias (putatively those with greater depressive symptoms) show greater remediation of response with stimulation. These are the two elements necessary for the development of closed-loop neuromodulation for mood disorders. With the demonstration of modulation of sensitive and quantifiable mood-related behavior, and with a neural target signature to track and predict modulation, this study may represent the first step toward a new future of adaptive closed-loop interventions for mood disorders targeting the dACC.

## Methods

### Participants

The study was approved by the Institutional Review Board at Baylor College of Medicine (IRB: H-18112). All participants provided written informed consent. Participants consisted of 16 subjects (6 female, mean age 38.25 years) with medically refractory epilepsy undergoing intracranial stereo EEG (sEEG) placement for intracranial epilepsy monitoring. sEEG implantation scheme was solely based on clinical criteria, with no influence from research considerations.

### Affective Bias Task (epilepsy patients)

Participants rated the emotional content of static, colorized photographs of adult human faces presented to a display monitor (Viewsonic VP150, 1920 x 1080) positioned at a distance of 57 cm. Faces consisted of emotional and neutral faces adapted from the NimStim Face Stimulus Set^43^. Happy, sad and neutral face exemplars (6 identities each; 3 male, 3 female) were morphed using a Delaunay tessellation matrix to generate subtle facial expressions ranging in emotional intensity from neutral to maximally expressive in steps of 10%, 30%, 50%, and 100% for happy and sad faces alike (Fig. 1a). The final stimulus set consisted of 54 stimulus exemplars (6 identities x 9 levels of intensity (100% sad, 50% sad, 30% sad, 10% sad, neutral, 10% happy, 30% happy, 50% happy and 100% happy)). For statistical modeling, trials were grouped into ‘subtle’ (neutral and 10%) and ‘overt’ (50% and 100%) separately for happy and sad faces, providing a total of four conditions. All stimuli were presented using the Psychtoolbox extensions for MATLAB^44^.

Trials began with the presentation of a white fixation cross presented on a black background for 1000 ms (jittered +/− 100 ms) followed by the simultaneous appearance of a face and the rating prompt (appearing on the left and right sides of the display, respectively; Fig. 1b). The rating prompt consisted of an active analog slider bar placed below text instructing patients to ‘Please rate the emotion.’ Participants used a computer mouse to indicate their rating by clicking a location on the slider bar. Ratings were recorded using a continuous scale ranging from 0 (‘Very Sad’) to 0.5 (‘Neutral’) to 1 (‘Very Happy’). Stimuli were presented in a blocked design in which all happy faces (plus neutral) appeared in one block while all sad faces (plus neutral) appeared in a separate block. Blocks consisted of one repetition of each image for a total of 30 randomized trials per block (6 identities x 5 levels of intensity). A recording session consisted of three blocks each of happy and sad faces, alternating between happy and sad (order counterbalanced across participants).

### Affective Bias Task (Amazon MTurk)

Stimuli and experimental design were like the task that was administered to epilepsy patients with a few exceptions. Given the nature of the task, we are unable to control the distance from or the size of the screen. We were also unable to add jitter between trials. Images were the same as the other study, and were presented in random order in blocked design (i.e. neutral plus sad faces in one block, neutral plus happy faces in a separated block). Four hundred MTurk participants were initially screened for depression symptoms, and the face rating task was administered to the participants exhibiting no initial symptoms via Qualtrics (n=86, Mean Age=38.5, M:F ratio=47:39). Averaged “norm” values were calculated for each face after administration of the task to the healthy controls through MTurk (the “expected score”) and then those values were subtracted from the ratings the patients in the Epilepsy Monitoring Unit (EMU) gave the same faces (the “observed score”) to determine each patient’s deviance from the norm value for each face (Observed-Expected).

### Affective Bias (stimulation experiment)

Stimuli were identical to those used in the main experiment. Experimental sessions began by applying brief 1-second pulses of stimulation during neurophysiological monitoring of the iEEG signal to ensure no presence of epileptic afterdischarge activity. Stimulation amplitude started at 0.5 mA and increased in 0.5 mA steps to 4 mA, and after not detecting any after-discharge activity, the experiment continued into the long-form stimulation phase. Biphasic symmetric rectangular-wave continuous stimulation was applied using a current-regulated device (CereStim R96, Blackrock Microsystems) at 130 Hz with a pulse width of 100 μsec while patients completed the Affective Bias task. Stimulation data was compared to non-stimulation sham runs of the task, which were collected just prior to the stimulation runs. Non-stimulation sham runs served as a baseline comparison. Patients were blinded from stimulation condition. One patient (‘YDX’) received bipolar stimulation, while the other 3 received monopolar stimulation.

### Intracranial EEG data collection

Neural signals were recorded from stereo EEG probes (Ad-Tech Medical Instrument Corporation, PMT Corporation) connected to a Cerebus data acquisition system (Blackrock Neurotech). All recording signals were amplified, filtered (high-pass 0.3 Hz first-order Butterworth, low-pass 500 Hz fourth-order Butterworth), and digitized at 2000 Hz. Trial onsets were marked using a photodiode which was placed in the lower right-hand corner of the visual display. Simultaneous with trial onsets, a white square (hidden from participant’s view) appeared at the location of the photodiode. The analog voltage response of the photodiode was recorded by the data acquisition system to ensure precise synchronization.

### Intracranial EEG data preprocessing

Based on visual inspection, channels with excessive noise artifacts, as well as channels containing ictal and interictal activity were excluded from data processing. Valid channels were then decimated to 500 Hz using a low-pass Chebyshev Type 1 IIR filter of order 8. Data were notch filtered to remove line noise (60 Hz, first harmonic, second harmonic) and then referenced according to a modified, single-dimension Laplacian approach, in which each electrode was referenced to the average of its adjacent neighbors along a sEEG probe. Contacts at either end of the probe were bipolar referenced to its immediate neighbor along the probe.

### Empirical mode oscillation detection (EMOD)

Selection of frequency bands for analysis has shifted away from using canonical frequency bands (i.e. theta, alpha, etc) in favor of data-driven approaches, in which a prominent feature is identified from the frequency domain (i.e. power spectral density information; PDS), and subsequent analyses focus activity at the identified feature^39,41,45^. Accurate estimation of the central frequency and magnitude of the feature requires removal of the 1/f aperiodic signal^45^. Existing 1/f removal methods rely on iterative model fitting/signal regression, which can introduce errors. The current report describes a novel approach to oscillatory feature detection that can isolate and remove the 1/f aperiodic signal without relying on model fitting. This method makes use of empirical mode decomposition (EMD), but instead of applying EMD to time-domain data, EMD is applied to frequency-domain data (i.e. PSD). EMD is an iterative process that results in n intrinsic mode functions (determined largely by the complexity of the input) and residue (or residual). Applied correctly, residue conforms to a strictly monotonic shape and is without any local extrema. For any data that conforms to the 1/f power law, residue is the linear 1/f function, which can simply be subtracted from the original input signal.

EMD is implemented using emdx.m^46^. After referencing, channel signal data are transformed from the time-domain into the frequency domain using a wavelet transformation (Morlet, 7 cycles per wavelet; frequencies were equally spaced on a logarithmic scale from 1 to 200 Hz).

This provides a channel-by-time matrix of power values for each data run. At each wavelet frequency we calculate the mean power across all data runs, which provides an estimate of power spectral density (PSD) for each channel. Once generated, the PSD is passed into an empirical mode decomposition algorithm (emdx.m)^46^. The output of EMD consists of a matrix of intrinsic mode functions (IMFs) as well as the residue. Channel feature profiles are created by either summing the IMFs, or by subtracting the residue from the original signal. In cases where the residue is not provided, it can be calculated by summing the IMFs and subtracting the summed IMFs from the original signal. Channel features can then be identified by searching feature profiles for significantly large local maxima. Two additional criteria were applied to our feature search: 1) we searched for local maxima in the range of 4-14 Hz, 2) we considered a feature to be valid only if it had a magnitude greater than 1 standard deviation. Channels were excluded from analysis if no significant feature was detected. Analysis frequency bands were created for each channel using a 4-Hz window centered around the peak frequency of the dominant detected feature.

### Intracranial EEG data analysis

After identifying a dominant feature for each channel, data were epoched using a time window beginning at stimulus onset and ending at the behavioral response. Trial averages were calculated as mean power across all time sample within the trial window, as well as across wavelet frequencies within the dominant frequency band. Condition averages were calculated separately for happy subtle, happy overt, sad subtle, and sad overt.

Data were then entered into a linear mixed effects model (lme4, citation), followed by t-tests using Satterthwaite-approximated degrees of freedom (emmeans). The dependent variable was power in the dominant feature frequency band, fixed effects of intensity (subtle and overt) and valence (happy and sad), and random effects of participant and electrode nested within participant. Separate models were run for each of five pre-defined ROIs: amygdala (AMY), dorsal anterior cingulate (dACC), dorsal anterior insula (daINS), ventral anterior insula (vaINS), and medial-orbital/ventral-medial prefrontal cortex (vmPFC).

### Trial-by-trial correlation analysis

To determine whether activity in any ROI was predictive of task behavior, trial-level emotional ratings were correlated with power in the dominant detected feature frequency band. This analysis was carried out separately for each recording channel. We then took an ROI based approach to test for statistical significance using one-sample t-tests. Correlation coefficients were first transformed using Fisher’s Z-transformation before being tested for significance using single-sample studentized t-tests. Three tests were run for each ROI: one in which all trials were included, a second in which only happy face trials were included, and a third in which only sad faces were included. We split all ROIs into left and right hemispheres except for the amygdala and vaINS, since there was an inadequate number of channels to complete split the ROIs into left and right hemispheres. Correlation coefficients (and corresponding ROI-level t-tests) were calculated separately for valence-coded ratings, intensity-coded ratings, and bias-coded ratings. In total, we ran 45 t-tests and corrected for multiple comparisons using a false discovery rate of 0.05. The resulting critical p-value for statistical significance was 0.0202. To calculate the average correlation at the level of the ROI, Fisher’s Z-transformed correlation coefficients were averaged before being converted back to Pearson rvalues using the inverse of Fisher’s Z-transformation.

### Electrode localization and ROI inclusion criteria

Freesurfer was used to align post-implantation CT brain scan from each patient, showing the location of the intracranial electrodes, to their preoperative structural T1 MRI scans. Electrode positions were manually marked using BioImage Suite 35. iELVis was used to overlay electrode location into the MRI. Electrodes were then assigned to an ROI based first on Freesurfer segmentation, which was then confirmed based on independent expert visual inspection. Electrodes were included if they were located within 5 mm of the grey matter boundary of an ROI.

The dorsal anterior cingulate was parceled according to^18,47^, in which the dACC extended from the vertical boundary placed orthogonal to the anterior end of the corpus callosum extending to the end boundary between the paracentral lobule and superior frontal gyrus. The anterior insula included gyri anterior of the central sulcus of the insula and was split into ventral and dorsal ROIs. Channels located inferior to the insula apex were assigned to the ventral ROI, channels superior to the apex were assigned to the dorsal ROI. All electrodes traversing the orbitofrontal and ventromedial cortex were assigned to the vmPFC ROI. For group level visualizations, electrode coordinates were linearly transformed into standard space (MNI305)^48^. From a total of 2472 channels, 423 channels were in an ROI. Of those significant neural features were detected in 228 channels: 19 in amygdala (left hemisphere = 6), 56 in dACC (left = 31), 56 in daINS (left = 25), 16 in vaINS (left = 8), and 39 in vmPFC (left = 16).

## Acknowledgements

The project received funding support from the National Institutes of Health (Bijanki: R01-MH127006, K01-MH116364, R21-A1-NS104953; Sheth: UH3-NS103549), the Caroline Wiess Law Fund for Research in Molecular Medicine, the ARCO Foundation, and the McNair Foundation.

## Author contributions

B.M., and K.B. designed research. B.M., P.K., M.M., J.A., R.M., J.G., V.K., L.L., A.M., B.S., and K.B. performed experiments. B.M., P.K., B.C., R.M., B.S., and Z.W. analyzed data. B.M., M.M., and K.B. wrote the paper. All authors discussed the results and contributed toward the manuscript.

---

**Table S1.** Trial-by-trial correlation comparisons.

| ROI | Hemi | Valence-coded ratings |  |  | Intensity-coded ratings |  |  | Bias-coded ratings |  |  |
| --- | --- | --- | --- | --- | --- | --- | --- | --- | --- | --- |
|  |  | *Rho* | *T-score* | *P-value* | *Rho* | *T-score* | *P-value* | *Rho* | *T-score* | *P-value* |
| AMY | Both | -0.07 | -2.84 | 0.011* | -0.05 | -3.36 | 0.004* | -0.01 | -0.38 | 0.706 |
|  | Right | -0.07 | -2.78 | 0.017* | -0.05 | -3.02 | 0.011* | -0.02 | -0.88 | 0.397 |
|  | Left | -0.06 | -1.05 | 0.343 | -0.05 | -1.48 | 0.200 | 0.02 | 0.80 | 0.459 |
| dACC | Both | 0.07 | 5.63 | <0.001* | -0.08 | -6.95 | <0.001* | 0.05 | 3.74 | <0.001* |
|  | Right | 0.05 | 2.72 | 0.012* | -0.10 | -5.78 | <0.001* | 0.04 | 1.82 | 0.081 |
|  | Left | 0.09 | 5.23 | <0.001* | -0.07 | -4.25 | 0.001* | 0.06 | 3.45 | 0.002* |
| daINS | Both | 0.04 | 2.08 | 0.042 | -0.06 | -4.99 | <0.001* | -0.01 | -0.51 | 0.615 |
|  | Right | 0.04 | 1.39 | 0.176 | -0.04 | -2.45 | 0.020* | 0.00 | -0.23 | 0.815 |
|  | Left | 0.04 | 1.59 | 0.124 | -0.08 | -4.91 | 0.001* | -0.01 | -0.49 | 0.626 |
| vaINS | Both | 0.01 | 0.47 | 0.649 | -0.06 | -2.33 | 0.034 | -0.04 | -1.39 | 0.186 |
|  | Right | -0.03 | -0.62 | 0.550 | -0.09 | -3.41 | 0.009* | -0.05 | -1.45 | 0.186 |
|  | Left | 0.06 | 1.69 | 0.142 | -0.02 | -0.41 | 0.697 | -0.01 | -0.33 | 0.752 |
| vmPFC | Both | 0.04 | 2.21 | 0.033 | -0.10 | -5.70 | <0.001* | 0.02 | 1.32 | 0.194 |
|  | Right | 0.02 | 0.93 | 0.362 | -0.13 | -4.84 | 0.001* | -0.01 | -0.48 | 0.637 |
|  | Left | 0.07 | 2.16 | 0.047 | -0.06 | -3.94 | 0.001* | 0.06 | 2.23 | 0.041 |

Statistical significance: \* < 0.05, FDR corrected

**Extended Data Fig. 1.**
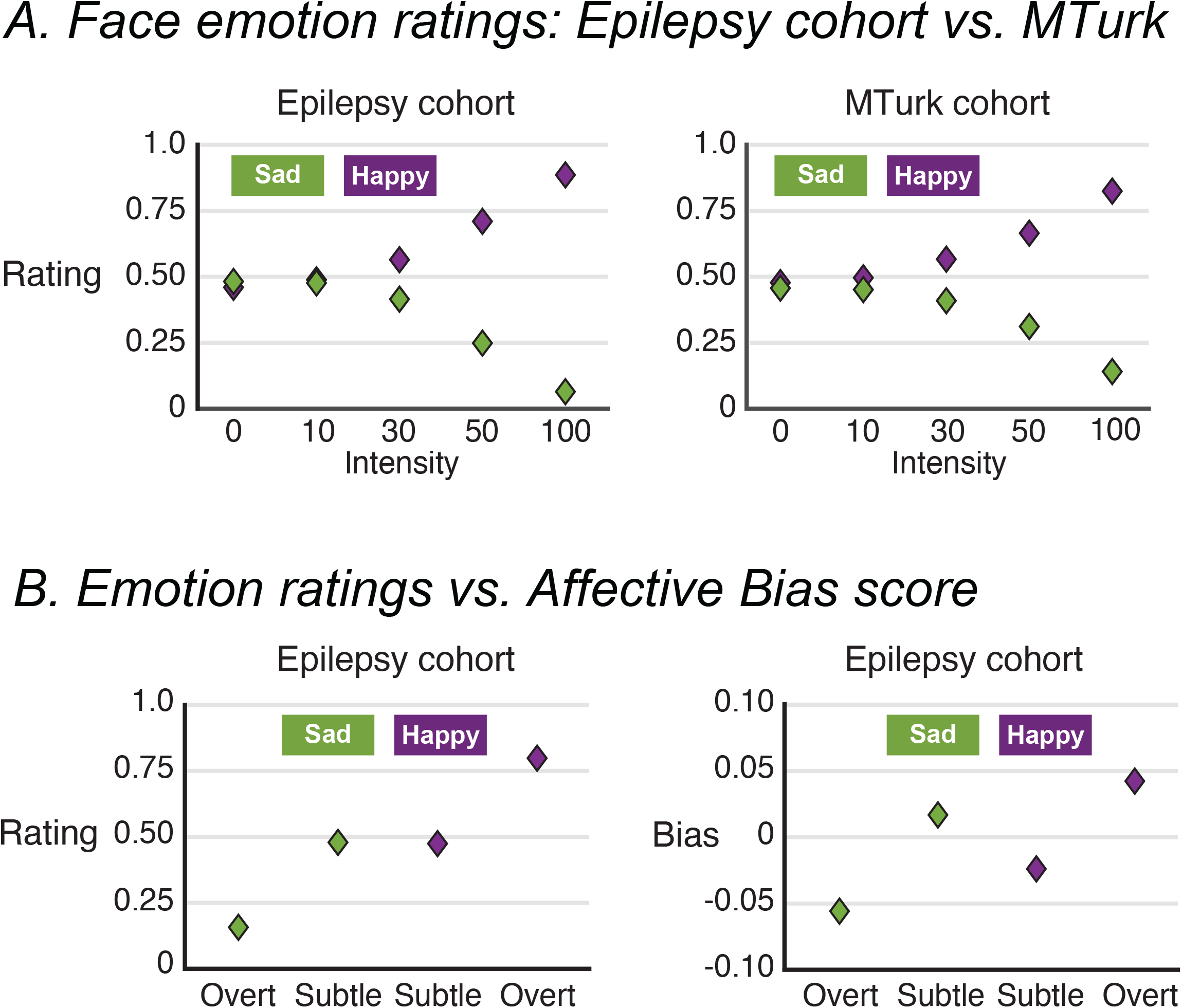
Affective Bias behavioral summary. **a** Average face emotion at each level of emotional intensity displayed separately for Happy (purple) and Sad (green) faces. Left panel: Epilepsy cohort (n = 16); right panel: Amazon MTurk cohort (n = 86). **b** Emotion ratings (left panel) compared against the affective bias scores (right panel) for the patients in the main iEEG study.: emotional ratings for each patient as a function of whether stimulation was off (x-axis) or on (y-axis). Dashed line represents line of equivalence. Ratings above the dashed line indicate ratings were higher while stimulation was on. **b** Right panel: difference between stimulation on and stimulation off (x-axis) compared against aver-

**Extended Data Fig. 2.**
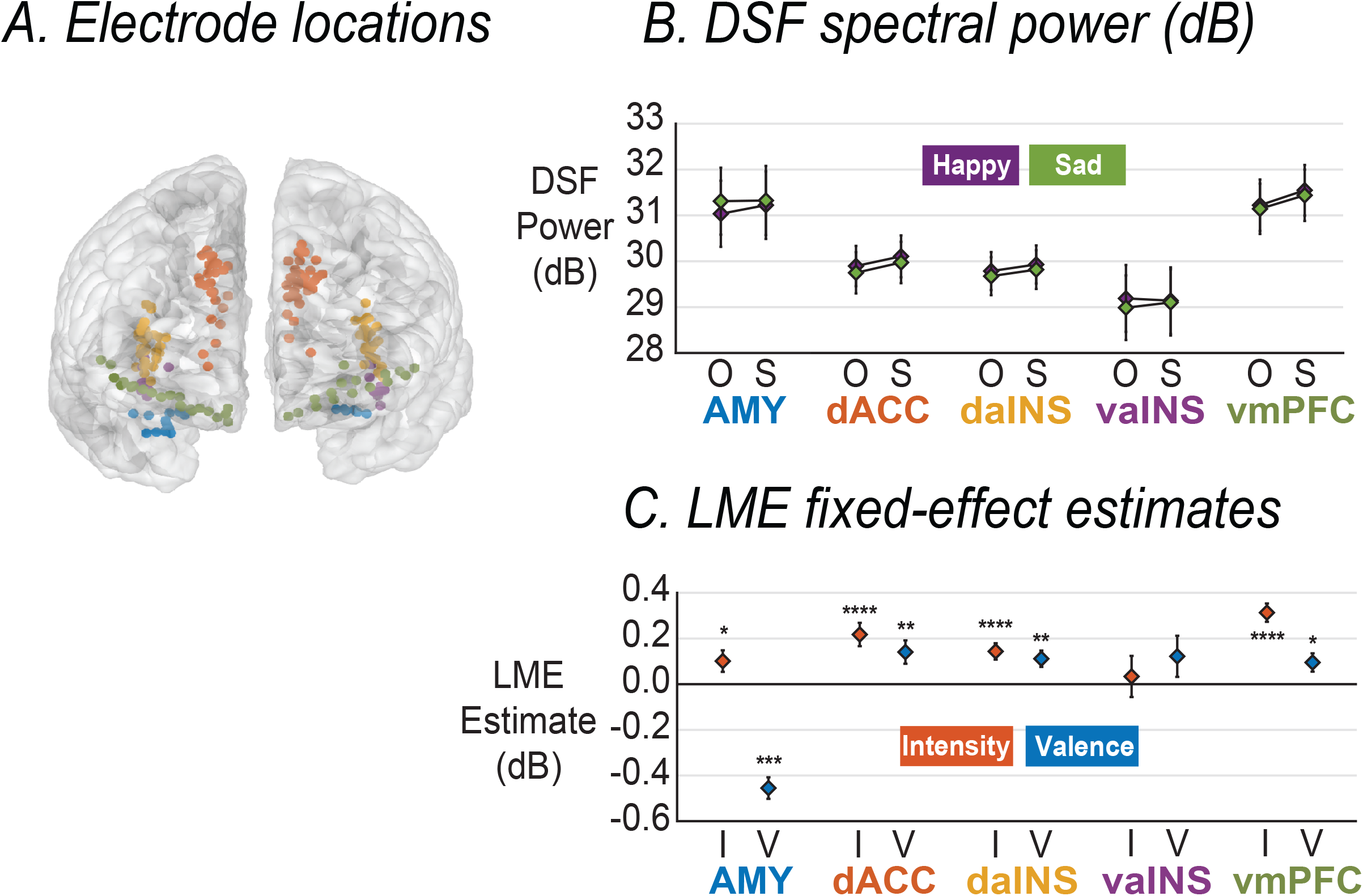
DSF spectral power: Intensity vs. Valence. **a** Electrode locations colored according to ROI. **b** Mean spectral power (dB) shown as a function of intensity (‘O’ = Overt, ‘S’ = Subtle) and valence (Happy vs. Sad) averaged across all channels for each ROI (error bars = +/− 1SEM). **c** Fixed-effect estimates (dB) for Intensity (I) and Valence (V) from a linear mixed-effects model run separately for each ROI (error bars = +/− 1 SEM; * = < 0.05, ** < 0.01, *** < 0.001, **** < 0.0001).

**Extended Data Fig. 3.**
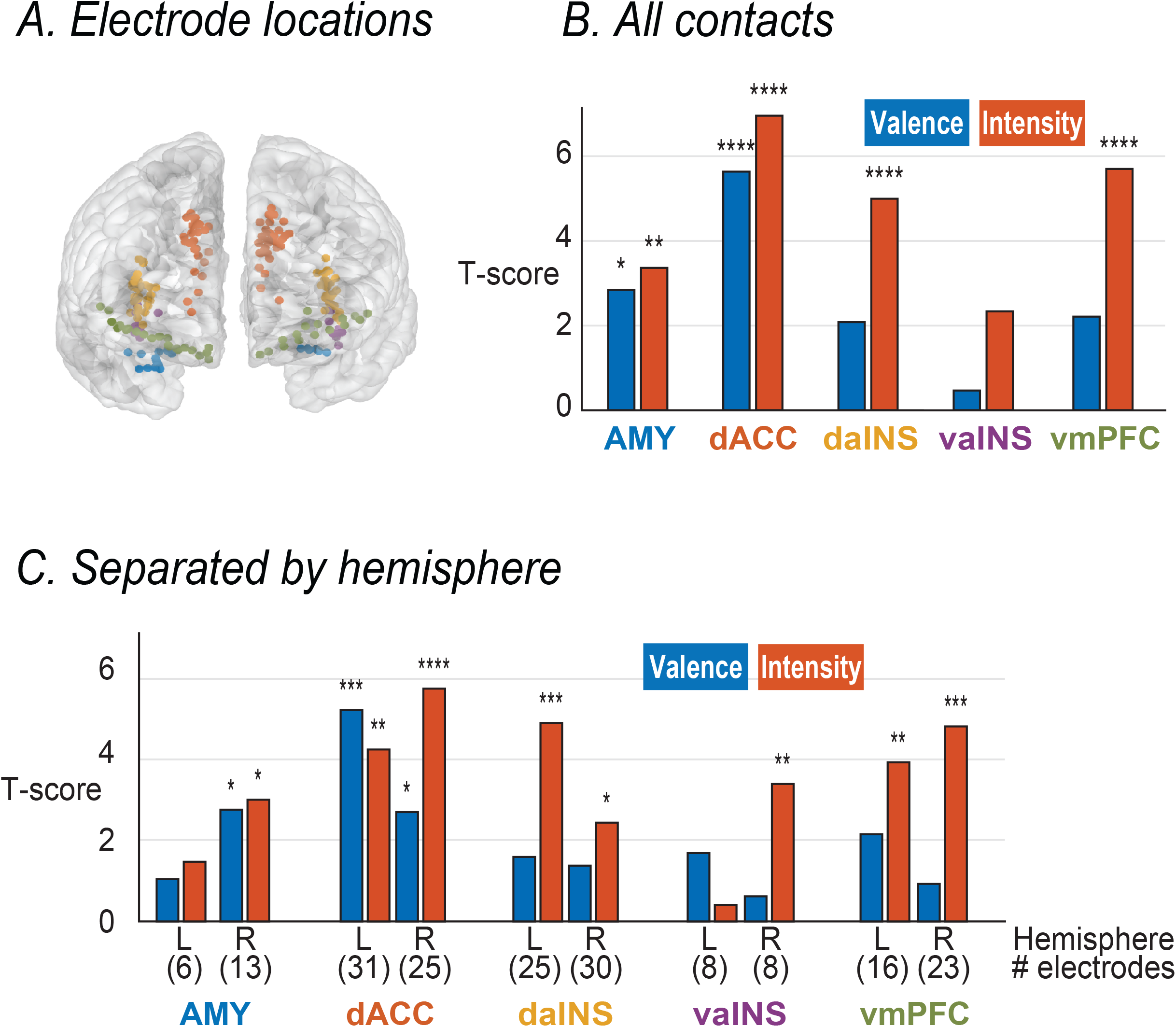
Predicting DSF spectral power from valence-coded vs. intensity-coded ratings. ROI-level comparisons of trial-by-trial correlations between spectral power and either intensity-coded ratings (orange bars) or valence-coded ratings (blue bars) displayed as a t-score for each ROI. **a** Electrode location colored according to ROI. **b** displays the results when all contacts are included regardless of hemisphere, **c** displays the results when analyses are run separately for left hemisphere (L) and right hemisphere contacts (R). Intensity-coded predictions are displayed as orange bars, Valence-coded predictions are displayed as blue bars (* = < 0.05, ** < 0.01, *** < 0.001, **** < 0.0001).

## References

1. Sabatinelli, D., Frank, D. W., & Filkowski, M. M. (2020). Emotional networks in the brain. In V. Zeigler-Hill & T. K. Shackelford (Eds.), Encyclopedia of personality and individual differences (pp. 1329–1338). Springer International Publishing. https://doi.org/10.1007/978-3-319-24612-3_511

2. Seeley, W. W. (2019). The salience network: A neural system for perceiving and responding to homeostatic demands. The Journal of Neuroscience, 39(50), 9878–9882. https://doi.org/10.1523/JNEUROSCI.1138-17.2019

3. Seeley, W. W., Menon, V., Schatzberg, A. F., Keller, J., Glover, G. H., Kenna, H., Reiss, A. L., & Greicius, M. D. (2007). Dissociable intrinsic connectivity networks for salience processing and executive control. The Journal of Neuroscience, 27(9), 2349–2356. https://doi.org/10.1523/JNEUROSCI.5587-06.2007

4. Adolphs, Ralph. (2002). Neural systems for recognizing emotion. Current Opinion in Neurobiology, 12(2), 169–177. https://doi.org/10.1016/S0959-4388(02)00301-X

5. Adolphs, R, Tranel, D., Damasio, H., & Damasio, A. (1994). Impaired recognition of emotion in facial expressions following bilateral damage to the human amygdala. Nature, 372(6507), 669–672. https://doi.org/10.1038/372669a0

6. Anders, S., Lotze, M., Erb, M., Grodd, W., & Birbaumer, N. (2004). Brain activity underlying emotional valence and arousal: a response-related fMRI study. Human Brain Mapping, 23(4), 200–209. https://doi.org/10.1002/hbm.20048

7. Menon, V., & Uddin, L. Q. (2010). Saliency, switching, attention and control: a network model of insula function. Brain Structure & Function, 214(5–6), 655–667. https://doi.org/10.1007/s00429-010-0262-0

8. Uddin, L. Q., Nomi, J. S., Hébert-Seropian, B., Ghaziri, J., & Boucher, O. (2017). Structure and function of the human insula. Journal of Clinical Neurophysiology, 34(4), 300–306. https://doi.org/10.1097/WNP.0000000000000377

9. Hiser, J., & Koenigs, M. (2018). The multifaceted role of the ventromedial prefrontal cortex in emotion, decision making, social cognition, and psychopathology. Biological Psychiatry, 83(8), 638–647. https://doi.org/10.1016/j.biopsych.2017.10.030

10. Giuliani, N. R., Drabant, E. M., & Gross, J. J. (2011). Anterior cingulate cortex volume and emotion regulation: is bigger better? Biological Psychology, 86(3), 379–382. https://doi.org/10.1016/j.biopsycho.2010.11.010

11. Piretti, L., Pappaianni, E., Gobbo, S., Rumiati, R. I., Job, R., & Grecucci, A. (2021). Dissociating the role of dACC and dlPFC for emotion appraisal and mood regulation using cathodal tDCS. Cognitive, Affective & Behavioral Neuroscience. https://doi.org/10.3758/s13415-021-00952-3

12. Yin, S., Liu, Y., Petro, N. M., Keil, A., & Ding, M. (2018). Amygdala Adaptation and Temporal Dynamics of the Salience Network in Conditioned Fear: A Single-Trial fMRI Study. ENeuro, 5(1). https://doi.org/10.1523/ENEURO.0445-17.2018

13. Manoliu, A., Meng, C., Brandl, F., Doll, A., Tahmasian, M., Scherr, M., Schwerthöffer, D., Zimmer, C., Förstl, H., Bäuml, J., Riedl, V., Wohlschläger, A. M., & Sorg, C. (2013). Insular dysfunction within the salience network is associated with severity of symptoms and aberrant inter-network connectivity in major depressive disorder. Frontiers in Human Neuroscience, 7, 930. https://doi.org/10.3389/fnhum.2013.00930

14. Williams, L. M. (2016). Precision psychiatry: a neural circuit taxonomy for depression and anxiety. The Lancet. Psychiatry, 3(5), 472–480.https://doi.org/10.1016/S2215-0366(15)00579-9

15. Mayberg, H. S., Lozano, A. M., Voon, V., McNeely, H. E., Seminowicz, D., Hamani, C., Schwalb, J. M., & Kennedy, S. H. (2005). Deep brain stimulation for treatment-resistant depression. Neuron, 45(5), 651–660. https://doi.org/10.1016/j.neuron.2005.02.014

16. Lozano, A. M., Mayberg, H. S., Giacobbe, P., Hamani, C., Craddock, R. C., & Kennedy, S. H. (2008). Subcallosal cingulate gyrus deep brain stimulation for treatment-resistant depression. Biological Psychiatry, 64(6), 461–467. https://doi.org/10.1016/j.biopsych.2008.05.034

17. Malone, D. A., Dougherty, D. D., Rezai, A. R., Carpenter, L. L., Friehs, G. M., Eskandar, E. N., Rauch, S. L., Rasmussen, S. A., Machado, A. G., Kubu, C. S., Tyrka, A. R., Price, L. H., Stypulkowski, P. H., Giftakis, J. E., Rise, M. T., Malloy, P. F., Salloway, S. P., & Greenberg, B. D. (2009). Deep brain stimulation of the ventral capsule/ventral striatum for treatment-resistant depression. Biological Psychiatry, 65(4), 267–275. https://doi.org/10.1016/j.biopsych.2008.08.029

18. Bijanki, K. R., Kovach, C. K., McCormick, L. M., Kawasaki, H., Dlouhy, B. J., Feinstein, J., Jones, R. D., & Howard, M. A. (2014). Case report: stimulation of the right amygdala induces transient changes in affective bias. Brain Stimulation, 7(5), 690–693. https://doi.org/10.1016/j.brs.2014.05.005

19. Drobisz, D., & Damborská, A. (2019). Deep brain stimulation targets for treating depression. Behavioural brain research, 359, 266–273. https://doi.org/10.1016/j.bbr.2018.11.004

20. Gur, R. C., Erwin, R. J., Gur, R. E., Zwil, A. S., Heimberg, C., & Kraemer, H. C. (1992). Facial emotion discrimination: II. Behavioral findings in depression. Psychiatry research, 42(3), 241–251. https://doi.org/10.1016/0165-1781(92)90116-k

21. Harmer, C. J., O’Sullivan, U., Favaron, E., Massey-Chase, R., Ayres, R., Reinecke, A., Goodwin, G. M., & Cowen, P. J. (2009). Effect of acute antidepressant administration on negative affective bias in depressed patients. The American Journal of Psychiatry, 166(10), 1178–1184. https://doi.org/10.1176/appi.ajp.2009.09020149

22. Surguladze, S. A., Young, A. W., Senior, C., Brébion, G., Travis, M. J., & Phillips, M. L. (2004). Recognition accuracy and response bias to happy and sad facial expressions in patients with major depression. Neuropsychology, 18(2), 212–218. https://doi.org/10.1037/0894-4105.18.2.212

23. Bijanki, K. R., Manns, J. R., Inman, C. S., Choi, K. S., Harati, S., Pedersen, N. P., Drane, D. L., Waters, A. C., Fasano, R. E., Mayberg, H. S., & Willie, J. T. (2019). Cingulum stimulation enhances positive affect and anxiolysis to facilitate awake craniotomy. The Journal of Clinical Investigation.

24. Dichter, G. S., Kozink, R. V., McClernon, F. J., & Smoski, M. J. (2012). Remitted major depression is characterized by reward network hyperactivation during reward anticipation and hypoactivation during reward outcomes. Journal of affective disorders, 136(3), 1126–1134. https://doi.org/10.1016/j.jad.2011.09048

25. Keedwell, P. A., Andrew, C., Williams, S. C., Brammer, M. J., & Phillips, M. L. (2005). The neural correlates of anhedonia in major depressive disorder. Biological psychiatry, 58(11), 843–853. https://doi.org/10.1016/j.biopsych.2005.05.019

26. Yee, D. M., Crawford, J. L., Lamichhane, B., & Braver, T. S. (2021). Dorsal Anterior Cingulate Cortex Encodes the Integrated Incentive Motivational Value of Cognitive Task Performance. The Journal of neuroscience: the official journal of the Society for Neuroscience, 41(16), 3707–3720. https://doi.org/10.1523/JNEUROSCI.2550-20.2021

27. Shenhav, A., Botvinick, M. M., & Cohen, J. D. (2013). The expected value of control: an integrative theory of anterior cingulate cortex function. Neuron, 79(2), 217–240. https://doi.org/10.1016/j.neuron.2013.07.007

28. Yin, Y., He, X., Xu, M., Hou, Z., Song, X., Sui, Y., Liu, Z., Jiang, W., Yue, Y., Zhang, Y., Liu, Y., & Yuan, Y. (2016). Structural and Functional Connectivity of Default Mode Network underlying the Cognitive Impairment in Late-onset Depression. Scientific reports, 6, 37617. https://doi.org/10.1038/srep37617

29. Jones, N. A., & Fox, N. A. (1992). Electroencephalogram asymmetry during emotionally evocative films and its relation to positive and negative affectivity. Brain and Cognition, 20(2), 280–299. https://doi.org/10.1016/0278-2626(92)90021-D

30. Schmidt, L. A., & Trainor, L. J. (2001). Frontal brain electrical activity (EEG) distinguishes valence and intensity of musical emotions. Cognition & Emotion, 15(4), 487–500. https://doi.org/10.1080/02699930126048

31. Gur, R. C., Skolnick, B. E., & Gur, R. E. (1994). Effects of emotional discrimination tasks on cerebral blood flow: regional activation and its relation to performance. Brain and cognition, 25(2), 271–286. https://doi.org/10.1006/brcg.1994.1036

32. Beraha, E., Eggers, J., Hindi Attar, C., Gutwinski, S., Schlagenhauf, F., Stoy, M., Sterzer, P., Kienast, T., Heinz, A., & Bermpohl, F. (2012). Hemispheric asymmetry for affective stimulus processing in healthy subjects--a fMRI study. Plos One, 7(10), e46931. https://doi.org/10.1371/journal.pone.0046931

33. Nimchinsky, E. A., Gilissen, E., Allman, J. M., Perl, D. P., Erwin, J. M., & Hof, P. R. (1999). A neuronal morphologic type unique to humans and great apes. Proceedings of the National Academy of Sciences of the United States of America, 96(9), 5268–5273. https://doi.org/10.1073/pnas.96.9.5268

34. Seeley, W. W., Carlin, D. A., Allman, J. M., Macedo, M. N., Bush, C., Miller, B. L., & Dearmond, S. J. (2006). Early frontotemporal dementia targets neurons unique to apes and humans. Annals of neurology, 60(6), 660–667. https://doi.org/10.1002/ana.21055

35. Shafir, R., Thiruchselvam, R., Suri, G., Gross, J. J., & Sheppes, G. (2016). Neural processing of emotional-intensity predicts emotion regulation choice. Social cognitive and affective neuroscience, 11(12), 1863–1871. https://doi.org/10.1093/scan/nsw114

36. Tamietto, M., Latini Corazzini, L., de Gelder, B., & Geminiani, G. (2006). Functional asymmetry and interhemispheric cooperation in the perception of emotions from facial expressions. Experimental brain research, 171(3), 389–404. https://doi.org/10.1007/s00221-005-0279-4

37. Omigie, D., Dellacherie, D., Hasboun, D., George, N., Clement, S., Baulac, M., Adam, C., & Samson, S. (2015). An Intracranial EEG Study of the Neural Dynamics of Musical Valence Processing. Cerebral cortex (New York, N.Y.: 1991), 25(11), 4038–4047. https://doi.org/10.1093/cercor/bhu118

38. Sonkusare, S., Nguyen, V. T., Moran, R., van der Meer, J., Ren, Y., Koussis, N., Dionisio, S., Breakspear, M., & Guo, C. (2020). Intracranial-EEG evidence for medial temporal pole driving amygdala activity induced by multi-modal emotional stimuli. Cortex; a journal devoted to the study of the nervous system and behavior, 130, 32–48. https://doi.org/10.1016/j.cortex.2020.05.018

39. Goyal, A., Miller, J., Qasim, S. E., Watrous, A. J., Zhang, H., Stein, J. M., Inman, C. S., Gross, R. E., Willie, J. T., Lega, B., Lin, J. J., Sharan, A., Wu, C., Sperling, M. R., Sheth, S. A., McKhann, G. M., Smith, E. H., Schevon, C., & Jacobs, J. (2020). Functionally distinct high and low theta oscillations in the human hippocampus. Nature communications, 11(1), 2469. https://doi.org/10.1038/s41467-020-15670-6

40. Donoghue, T., Schaworonkow, N., & Voytek, B. (2021). Methodological considerations for studying neural oscillations. The European journal of neuroscience, 10.1111/ejn.15361. Advance online publication. https://doi.org/10.1111/ejn.15361

41. Miller, J., Watrous, A. J., Tsitsiklis, M., Lee, S. A., Sheth, S. A., Schevon, C. A., Smith, E. H., Sperling, M. R., Sharan, A., Asadi-Pooya, A. A., Worrell, G. A., Meisenhelter, S., Inman, C. S., Davis, K. A., Lega, B., Wanda, P. A., Das, S. R., Stein, J. M., Gorniak, R., & Jacobs, J. (2018). Lateralized hippocampal oscillations underlie distinct aspects of human spatial memory and navigation. Nature communications, 9(1), 2423. https://doi.org/10.1038/s41467-018-04847-9

42. Watrous, A. J., Miller, J., Qasim, S. E., Fried, I., & Jacobs, J. (2018). Phase-tuned neuronal firing encodes human contextual representations for navigational goals. eLife, 7, e32554. https://doi.org/10.7554/eLife.32554

43. Tottenham, N., Tanaka, J. W., Leon, A. C., McCarry, T., Nurse, M., Hare, T. A., Marcus, D. J., Westerlund, A., Casey, B. J., & Nelson, C. (2009). The NimStim set of facial expressions: judgments from untrained research participants. Psychiatry research, 168(3), 242–249. https://doi.org/10.1016/j.psychres.2008.05.006

44. Brainard D. H. (1997). The Psychophysics Toolbox. Spatial vision, 10(4), 433–436.

45. Donoghue, T., Haller, M., Peterson, E. J., Varma, P., Sebastian, P., Gao, R., Noto, T., Lara, A. H., Wallis, J. D., Knight, R. T., Shestyuk, A., & Voytek, B. (2020). Parameterizing neural power spectra into periodic and aperiodic components. Nature neuroscience, 23(12), 1655–1665. https://doi.org/10.1038/s41593-020-00744-x

46. Cohen, S., & Collins, M. (2014). A Provably Correct Learning Algorithm for Latent-Variable PCFGs. In Proceedings of the 52nd Annual Meeting of the Association for Computational Linguistics (pp. 1052–1061). Association for Computational Linguistics. http://aclweb.org/anthology/P14-1099

47. McCormick, L. M., Keel, P. K., Brumm, M. C., Bowers, W., Swayze, V., Andersen, A., & Andreasen, N. (2008). Implications of starvation-induced change in right dorsal anterior cingulate volume in anorexia nervosa. The International journal of eating disorders, 41(7), 602–610. https://doi.org/10.1002/eat.20549

48. Magnotti, J. F., Wang, Z., & Beauchamp, M. S. (2020). RAVE: Comprehensive opensource software for reproducible analysis and visualization of intracranial EEG data. NeuroImage, 223, 117341. https://doi.org/10.1016/j.neuroimage.2020.117341

